# Guiding and interpreting brain network classification with transcriptional data

**DOI:** 10.1101/2020.05.15.099028

**Authors:** Mengbo Li, Daniel Kessler, Jesús Arroyo, Saskia Freytag, Melanie Bahlo, Elizaveta Levina, Jean Yang

## Abstract

The investigation of brain networks has yielded many insights that have helped to characterise many neurological and psychiatric disorders. In particular, network classification of functional magnetic resonance imaging (fMRI) data is an important tool for the identification of prognostic and diagnostic biomarkers of brain connectivity disorders such as schizophrenia and depression. However, existing generic network classification methods provide no direct information on the underlying molecular mechanisms of the selected functional connectivity features when applied to fMRI data. To address this, we propose a novel fMRI network classification method that incorporates brain transcriptional data using a user-specified gene set collection (GSC) to construct feature groups for use in classification of brain connectivity data. The use of GSCs are an opportunity to incorporate knowledge of potential molecular mechanisms which may be associated with a disease. The inclusion of transcriptional data yields improved prediction accuracy on publicly available schizophrenia fMRI data for several of the GSCs we consider. We also introduce a post-hoc interpretation framework to provide transcriptional-data-guided biological interpretations for discriminative functional connectivity features identified by existing fMRI network classification methods.

## Introduction

Network analysis of functional brain imaging data has been widely applied in studies of neurological disorders and has facilitated the discovery of functional biomarkers in many complex brain diseases [6, 10, 39], but the molecular and neurobiological mechanisms underlying brain functional connectivity patterns remain elusive [36, 37]. Previous studies [23, 37, 43] on neuroimaging network classification in mental disorders that provided feature selection typically sought to biologically interpret algorithmically selected functional connectivity features post hoc, based on available literature on the known functions of certain brain regions. At the same time, it is known that differential transcription across distinct brain regions has an impact on brain functions [20, 30], raising the question of how changes in gene expression associated with a mental disorder may be connected to disrupted brain functional organisation (note that in this paper we sometimes use the phrase “gene expression” when discussing transcriptional data). Recent work [30, 34] analysed transcriptional networks that focused on gene-gene correlations across distinct brain regions, and gene ontology enrichment analyses were used to assess the functional relevance of identified gene modules, but these studies did not examine the associated changes in functional connectivity patterns.

Traditionally, integrative analysis on multimodal data requires measurements of different modalities from the same cohort of subjects. For instance, genome-wide association studies (GWAS) on brain imaging phenotypes require subject-matched structural or functional magnetic resonance imaging (fMRI) scans and genome-wide genetic data. However, functional imaging and antemortem transcriptional data are highly unlikely to be available from the same brain at the same time as expression assays require invasive access to brain tissues [2]. A recent paper [12] reviewed common approaches to link the functional connectome with the whole brain transcriptome from the Allen Human Brain Atlas (AHBA), an anatomically comprehensive human brain transcriptional atlas [19]. They include testing for correlations between measurements of brain imaging based disease phenotypes and (i) the spatial transcriptional profiles of a particular gene, (ii) eigengenes of disease associated pathways or gene ontology terms [4], or (iii) the spatial expression patterns of known risk genes. Another approach is to perform gene set enrichment tests on genes previously found to be associated with normative imaging phenotypes of the disease since many genes are functionally related [12]. Although these methods only provide a list of priority genes for further study, there is evidence that spatial variations in brain transcriptional profiles closely track variations in functional organisation of the brain [12].

Here, we introduce a transcriptional-data-guided functional network classification method to enable biologically annotated feature selection in functional brain network classification via the regional matching of multimodal data. Our case study on the publicly available data set shows that transcriptional-data-guided classification of functional connectivity facilitates the identification of underpinning molecular mechanisms for disrupted functional organisations in complex neurological diseases.

## Materials and Methods

### Data sets

Processed resting state fMRI networks for subjects (54 schizophrenic and 70 healthy control) from the Center for Biomedical Research Excellence (COBRE) schizophrenia fMRI data [1, 41] were downloaded from the `graphClass` R package [3]. Whole brain microarray gene expression data were downloaded from the Allen Human Brain Atlas (AHBA) at http://human.brain-map.org/static/download?rw=t. Both COBRE fMRI data and AHBA microarray gene expression samples were registered to the MNI152 1mm space during initial data pre-processing. Gene set information was downloaded from the Molecular Signatures Database (MSigDB, v7.0) [26]. Gene-schizophrenia association scores were obtained from DiGSeE [24], a MEDLINE abstract text mining search engine.

### Data processing

Regions of interest (ROIs) from the Power et al. brain parcellation [35] were used as nodes to construct the brain networks. Time course data were extracted from the ROIs defined by the Power parcellation. The 1, 488 AHBA gene expression samples were also mapped to the same brain parcellation and assigned to ROIs. Out of the 264 ROIs in the Power parcellation, 248 had at least one AHBA sample mapped to it. The remaining 16 ROIs which had no AHBA samples mapped to them were excluded from our transcriptional-data-guided method, but retained for use in other methods that do not directly depend upon transcriptional data (e.g., the baseline network classifiers and the post-hoc interpretation framework proposed below). ROI 75 was also discarded as it is missing in COBRE data. In summary, 247 ROIs with mapped transcriptional profiles were used as nodes to construct each transcriptional brain network. Details on the AHBA brain tissue sample to Power ROI mapping, as well as other gene expression data processing steps, are provided in Supplementary Methods.

### brainClass : Transcriptional-data-guided brain network classification

The proposed classification method has two major components, both with several steps.

#### (A) Construction of Gene Set Edge Groups (Figure 1a)

The transcriptional data is first partitioned by a *gene set collection (GSC)* under consideration. A weighted, undirected ROI-wise transcriptional network is then calculated for each gene set from the GSC as follows. In the transcriptional network of gene set *x*, the weight 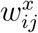 of an edge (*i, j*) between ROIs *i* ≠ *j* is computed as the absolute value of the Pearson correlation coefficient of the gene expression levels of genes in set *x* between ROI *i* and ROI *j*. Next, a very stringent hard-thresholding criterion is applied to obtain a subset of the highest-weighted edges in each network, defined by

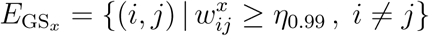

where *η*_0.99_ is the 99th percentile of all edge weights in this network.

**Figure 1:**
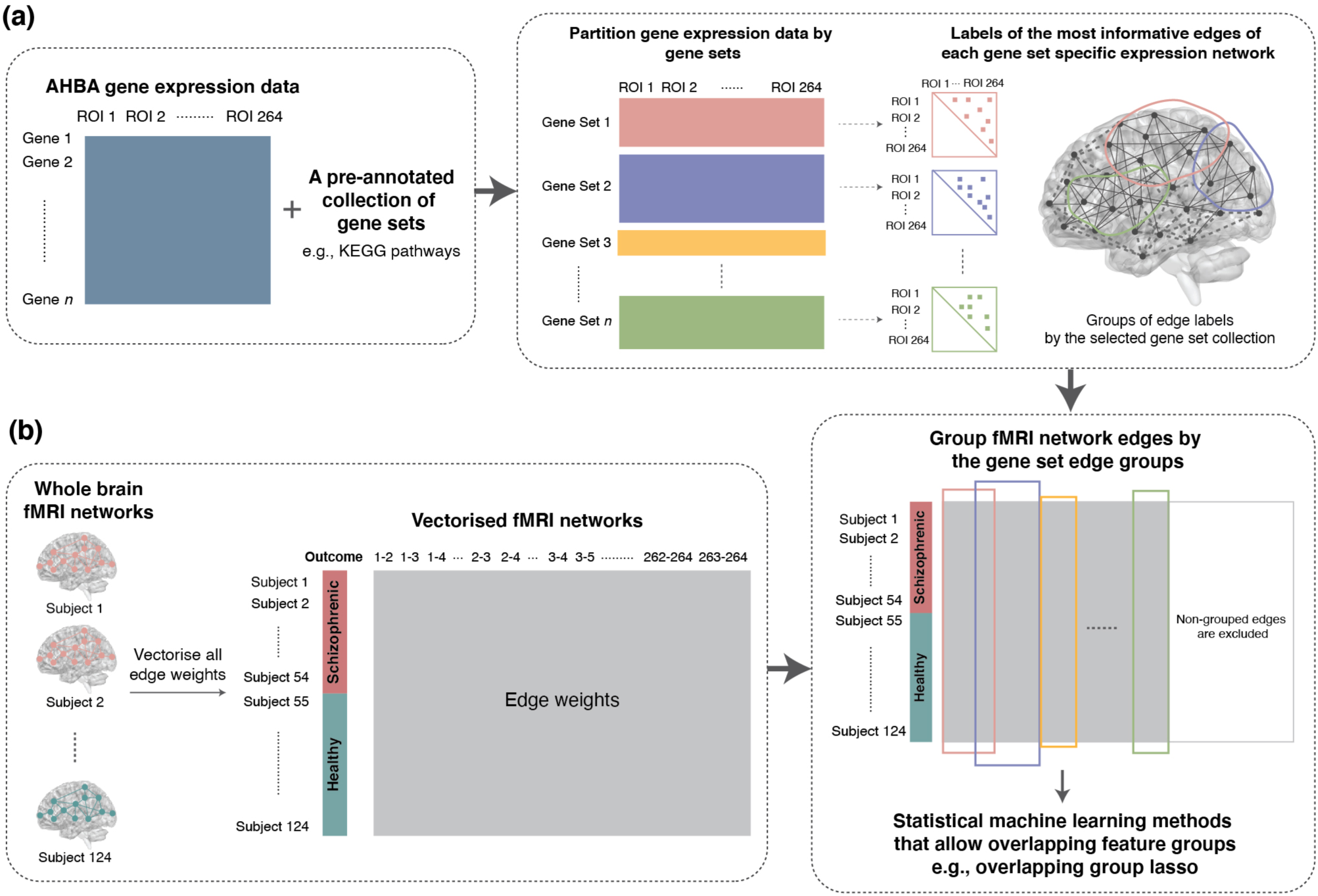
Schematic illustration of `brainClass`. (a) Construction of gene set edge groups based on transcriptional data. In the left panel, processed transcriptional data are represented as a data matrix with rows corresponding to genes and columns to ROIs. Given a GSC of interest, the transcriptional data matrix is partitioned by each gene set, and the ROI-wise transcriptional network is calculated for each gene set. Labels of the most highly weighted edges in each gene set expression network form a gene set edge group, labelled by the name of the corresponding gene set. (b) fMRI network classification. Functional connectivity edge weights are used as the features, which are grouped according to the previously constructed gene set edge groups. (ROI: Region of interest; GSC: gene set collection.)

#### (B) fMRI Network Classification (Figure 1b)

For each subject, the fMRI network is constructed using the Power ROIs as nodes. The Pearson correlation coefficient of the fMRI time courses between ROIs is used as the connectivity measure between nodes. Edge weights of each fMRI network are treated as a long feature vector in classification. See Supplementary Methods for details regarding fMRI data.

For fMRI network classification, we make use of statistical classification methods that perform feature selection among overlapping feature groups. Functional edges are grouped according to the gene set edge groups constructed in component (A) to form (potentially overlapping) feature groups, and edges that do not belong to any feature group are excluded from training. The current default implementation of `brainClass` uses logistic regression with the overlapping group lasso penalty implemented in the `grpregOverlap`R package [42], which allows user-defined overlapping feature groups.

#### Gene Set Prioritisation

With feature selection results from classification, gene sets within a given GSC can be ranked by any user-defined *gene set prioritisation score* to quantify their relevance to functional connectivity. In the current implementation, we utilise a measure for overlap defined as

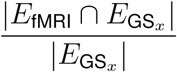

for each gene set *x*, where |·| denotes the cardinality of a set, and *E*_fMRI_ is defined as the set of edges selected (with a non-zero fitted coefficient) in each fold of a 10-fold cross-validation. Gene sets ranked in top 5% by the above measure in all ten folds in all 50 repetitions of 10-fold cross-validations are considered prioritised in the GSC of interest.

### Evaluation of brainClass

*Selection of GSC* The performance of `brainClass` was evaluated when various types of GSCs were used for the construction of gene set edge groups. The proposed algorithm was also evaluated when non-informative gene sets were utilised, as discussed below. Non-informative GSCs were generated with 606 gene sets each, which is the median size of all evaluated biologically meaningful GSCs (Figure 2c). The size of each individual gene set was randomly drawn from the distribution of all gene set sizes shown in Figure 2c, regardless of the category.

**Figure 2:**
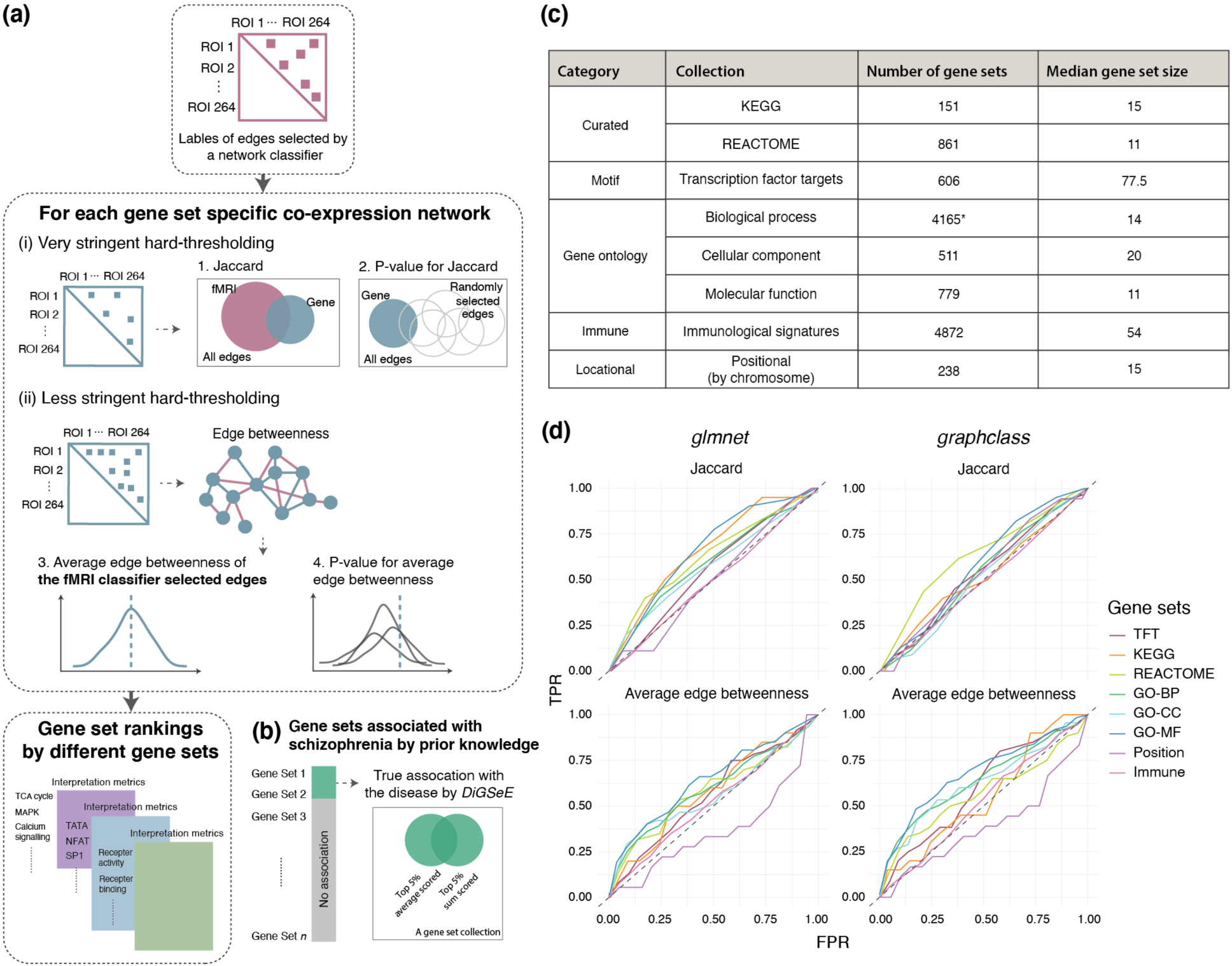
Post-hoc interpretation framework applied to existing fMRI network classification methods. (a) Schematic illustration of the post-hoc interpretation framework. (b) Gene sets with reported associations with schizophrenia in MEDLINE by DiGSeE. (c) Summary statistics on the pre-processed GSCs. Note: when evaluating the performance of the transcriptional-data-guided fMRI-network classifier with the GO GSC, we filtered to include only gene sets with size between 10 and 30, resulting in 1,492 gene sets. (d) Receiver operating characteristic (ROC) curves for two post-hoc interpretation metrics versus constructed DiGSeE scores. (GSC: gene set collection; TPR: true positive rate; FPR: false positive rate; TFT: transcription factor target; GO: Gene Ontology; BP: biological process; CC: cellular component; MF: molecular function.)

### Post-hoc interpretation framework

We propose a framework for the post-hoc interpretation of an edge set selected by a classifier (see Figure 2 for a schematic overview). Let *E*_fMRI_ denote the set of edges selected by an fMRI network classifier, consisting of edges with a non-zero estimated coefficient. As a post-hoc interpretation metric, we can compare the edge sets *E*_fMRI_ and 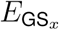, the subset of highest-weight edges in the transcriptional connectivity network of gene set *x*. We measure their similarity by the Jaccard index, defined as

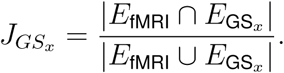

We then repeatedly select *n* = |*E*_fMRI_| edges at random from the set of all possible edges to obtain a null distribution for the Jaccard index and an estimated *P* -value for 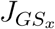.

Another metric is devised based on the edge betweenness centrality, defined as the number of the shortest paths via the edge in a network [16]. Because the strict threshold we previously applied resulted in highly disconnected graphs, we instead apply a less stringent threshold to obtain a binary transcriptional network of gene set *x*:

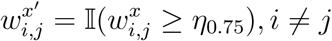

where *η*_0.75_ is the 3*rd* quartile of all edge weights **w**^*x*^, retaining 25% of edges with the highest weights instead of 1% as we did before. For this network, we calculate the average edge betweenness centrality for the edges that also belong to *E*_fMRI_, for each gene set *x*. We then generate a null distribution by randomly choosing *n* = |*E*_fMRI_| edges to serve as the pseudo *E*_fMRI_ set and repeat the above procedure: we intersect each 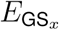 with the pseudo *E*_fMRI_ set and calculate the average edge betweenness centrality of these edges. We use this null distribution to obtain an estimated *P* -value for each gene set. We use the Jaccard index, the average edge betweenness, and the *P* -values associated with each to rank gene sets. The same approach is applicable to any other user-defined interpretation metric.

### Application of the post-hoc interpretation framework to COBRE data

#### Feature selection results evaluated by the post-hoc interpretation framework

We compared the functional edge selection results of our method to the two existing fMRI network classification methods we use as benchmarks. For logistic regression with the ordinary lasso penalty, `glmnet` [14], *E*_fMRI_ was defined as the set of edges selected (with a non-zero fitted coefficient) in at least one fold of at least ten of the 50 repetitions of 10-fold cross validation. For the overlapping lasso penalty in `graphClass` [3], as recommended by the authors, stability selection [27] was used to calculate variable selection probabilities, by randomly splitting the data and calculating the percentage of times across all random splits that each variable was chosen. Edges chosen more than half the time were selected.

#### Evaluation procedure

The effectiveness of different GSCs in explaining the functional edge selection results by other existing fMRI network classifiers were evaluated. Schizophrenia relevance scores were constructed for each gene set based on the gene-schizophrenia association scores extracted from DiGSeE [24] and log-transformed. We consider both (i) the sum of gene-based disease association scores (for identifying gene sets driven by a small number of strong effects) and (ii) the average disease association score for each set. Within each GSC, gene sets with a top 5% *sum score* or a top 5% *average score* were categorised as the true positives, or in other words, to be associated with schizophrenia, and true negatives otherwise (Figure 2b). The accuracy of gene set categorisation by each interpretation metric was evaluated by the receiver operating characteristic (ROC) curve constructed with a step size of 5%, where the true positive rate is plotted against the false positive rate, and the corresponding area under the curve (AUC). This procedure was repeated to obtain a distinct AUC score for each combination of GSC and classifier.

## Results

### brainClass: A new method for brain network classification guided by transcriptional data

Our novel fMRI network classification method, `brainClass`, uses gene expression across brain regions to guide connectivity-based feature selection, with the goal of bridging the gap between the functional biomarkers of neurological disorders and their underpinning molecular mechanisms. Firstly, brain regional matching is performed between whole brain transcriptional data and fMRI data via the spatial coordinates of each sample with the aid of a standardised brain template (e.g., MNI152 1mm). Both modalities of data are then mapped into a selected brain parcellation, where the pre-defined ROIs are used as nodes to construct both transcriptional and fMRI networks. Next, feature groups are strategically constructed and annotated to incorporate the biologically relevant gene set information, such as pre-defined biological pathways or gene ontology (GO) terms [4]. Finally, we perform network classification using statistical classification methods that allow overlapping feature groups.

Application on COBRE data shows the transcriptional-data-guided fMRI network classifier outperformed the baseline methods in terms of the classification accuracy when guided by certain GSCs (Figure 3a). Compared with `glmnet` (82%), `brainClass` implemented with any tested GSCs provided higher classification accuracy except for the positional gene sets (80%). The average cross-validation prediction accuracies of `brainClass` with feature groups constructed via GO molecular function terms (92%) and filtered GO biological process (only gene sets sized between 10 and 30 were used [22]) terms (94%) were also higher than that of `graphClass` (91%).

**Figure 3:**
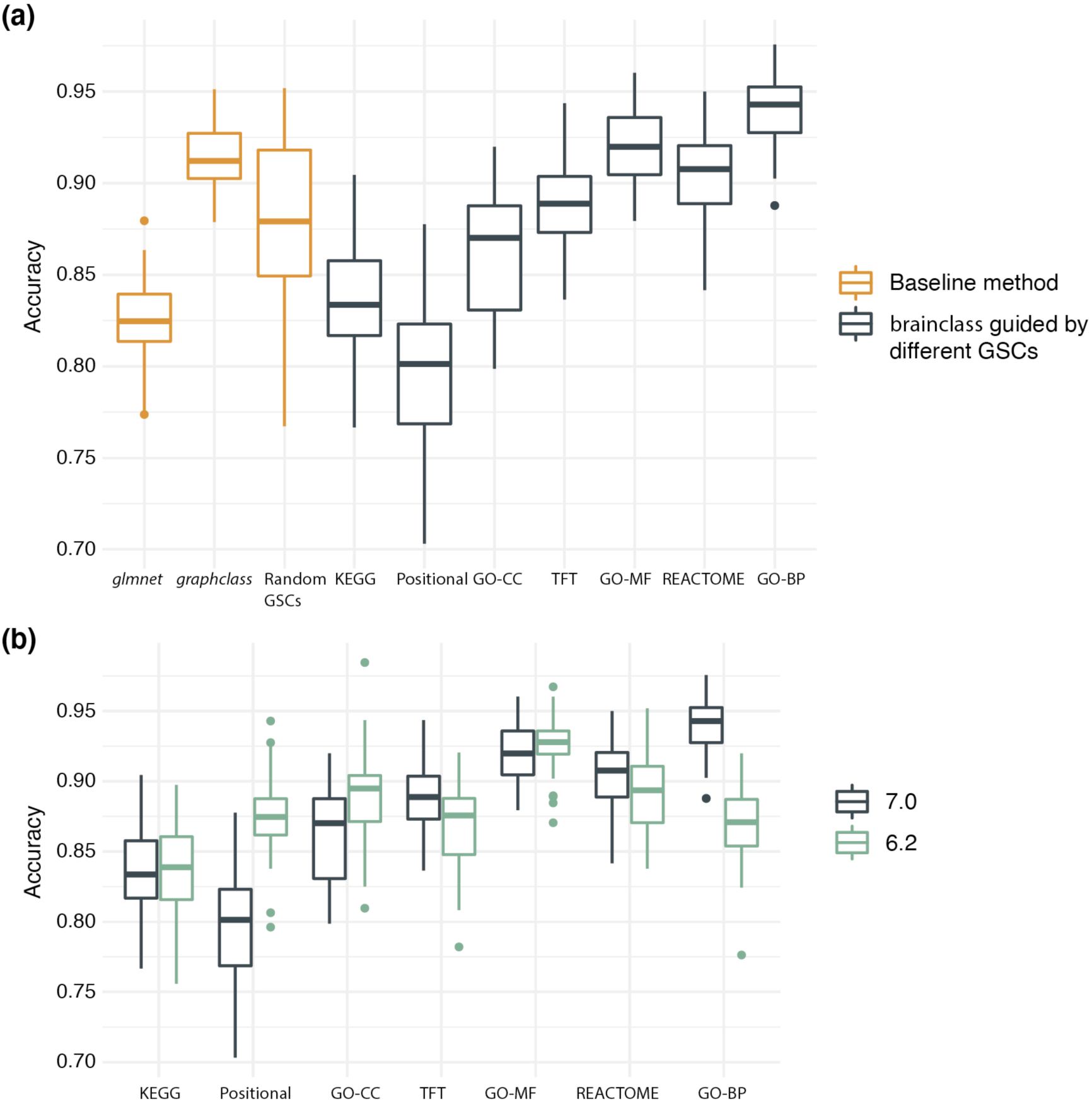
Evaluation of `brainClass`. (a) Prediction accuracy of the gene-expression guided fMRI network classification algorithm compared with baseline methods. We evaluated all fMRI network classifiers on the COBRE schizophrenia fMRI data in terms of the prediction accuracy in discriminating schizophrenic patients (*n* = 54) from healthy controls (*n* = 70) guided by different GSCs (obtained from MSigDB v7.0). Accuracy was calculated as the average prediction accuracy of 10-fold cross-validations in 50 repeats. (b) Prediction accuracy of `brainClass` guided by the gene set information provided in versions v6.2 and v7.0 of MSigDB.

### Classification performance depends on the GSC utilised in feature group construction

Prediction accuracy varied when different GSCs were used, as expected, since classification results depend strongly on feature groups determined by the GSC. Compared with using randomly generated GSCs, Figure 3a shows that the transcription factor targets (TFTs), GO molecular function terms, REACTOME pathways and filtered GO biological process terms outperformed the random GSCs (median accuracies of 89%, 92%, 91% and 94% respectively versus 88%; see the figure for a measure of CV variation around these median accuracies, represented by boxplots).

We observed reasonably high network classification accuracy with a small number of features selected by the baseline classifier of `glmnet` on the COBRE data. The median classification accuracy over 50 repetitions of 10-fold cross-validation was 82% (Figure 3a) for the ordinary lasso via `glmnet`. The number of unique features selected across 10 folds of each cross-validation repetition ranges from 200 to 300, out of the total of 34, 453 (263 choose 2) features, or below 1% on average. On the contrary, for `graphClass`, almost all features were always selected, to obtain the median estimation of classification accuracy of 91% (Figure 3a).

In `brainClass`, the GSC used in feature group construction influences the number of selected features by the proposed method. For example, 14, 721 functional edges were included in training by the positional GSC and 7, 057 edges by the GSC of TFT. In our analysis, the regularisation parameter λ `grpregOverlap` of was tuned to 5 × 10^−3^ so that the implemented algorithm selected around 10% features that had non-zero coefficients. Although sparse models (with few features selected) commonly have lower prediction accuracy in high-dimensional settings [18], the median prediction accuracy over repeated cross-validation was considerably higher when the proposed classifier was guided by TFT (89%) than by the positional GSC (80%). This implies that strategic construction of feature groups can potentially improve prediction accuracy and promote sparsity at the same time. The numbers of features included in training and the distribution of the number of selected features by different GSCs are provided in Supplementary Results.

### Quality of biological information affects prediction accuracy of brainClass

As curated gene set information evolves, the performance of the transcriptional-data-guided fMRI network classification algorithm can also improve. The prediction accuracy of `brainClass` was also evaluated when GSC information was downloaded from the previous release of MSigDB, v6.2 (Figure 3b). The current version (v7.0) implemented a major overhaul in GSCs including the positional gene sets and GO term GSCs compared to the older v6.2. There was also a major update in REACTOME pathways from v6.2 to v7.0, for which the prediction accuracy was improved when the fMRI network classification was guided by the updated pathway information (from 89% to 91%). An even bigger improvement was observed when GO biological process terms (filtered for gene sets sized between 10 and 30 in both versions) were utilised for feature group construction in classification (from 87% to 94%). The positional GSC yielded relatively poor performance (87% in v6.2 and 80% in v7.0), which is not surprising as we expect limited biologically relevant implications on schizophrenia by the physical location of genes on each chromosome.

### Selected features have an inherent biological interpretation

A major advantage of `brainClass` is the direct connection between selected connectivity features and their potential molecular underpinnings. By directly comparing the set of selected functional edges with the most highly weighted transcriptional edges in each gene set expression network, gene sets can be ranked by the proposed gene set prioritisation metric. For example, Table 1 shows the gene sets ranked in the top 5% by the implemented gene set prioritisation metric in each fold of the 50 repeats of 10-fold cross-validations when `brainClass` was guided by REAC-TOME pathways. Note that the DiGSeE gene set scores are not directly used by the `brainClass` classification method, but displayed here for reference only.

**Table 1:** REACTOME pathways with a prioritisation score in top 5% in all ten folds in all 50 repeats of 10-fold cross-validations, shown in alphabetical order together with the constructed DiGSeE gene set schizophrenia relevance scores of each pathway.

| Gene set | Sum score | Average score |
| --- | --- | --- |
| Adaptive Immune System | 7.04 | 0.04 |
| Asparagine N-Linked Glycosylation | 2.59 | 0.04 |
| Axon Guidance | 35.87 | 0.19 |
| Cell Cell Communication | 3.74 | 0.07 |
| Cell Cycle | 8.05 | 0.06 |
| Cellular Responses to External Stimuli | 10.39 | 0.09 |
| Cellular Responses to Stress | 9.06 | 0.09 |
| Cytokine Signaling in Immune System | 54.84 | 0.26 |
| Developmental Biology | 48.26 | 0.16 |
| Disease | 56.25 | 0.22 |
| Gene Expression Transcription | 35.95 | 0.13 |
| Generic Transcription Pathway | 35.95 | 0.15 |
| Hemostasis | 27.03 | 0.14 |
| Innate Immune System | 38.95 | 0.15 |
| Integration of Energy Metabolism | 7.29 | 0.19 |
| Intracellular Signaling by Second Messengers | 26.41 | 0.31 |
| MAPK Family Signaling Cascades | 31.12 | 0.34 |
| Metabolism of Amino Acids and Derivatives | 21.20 | 0.21 |
| Metabolism of Carbohydrates | 1.31 | 0.02 |
| Metabolism of Lipids | 22.82 | 0.12 |
| Metabolism of RNA | 1.04 | 0.01 |
| Neuronal System | 73.54 | 0.40 |
| Other Interleukin Signaling | 30.75 | 0.34 |
| Phospholipid Metabolism | 3.83 | 0.06 |
| Platelet Activation Signaling and Aggregation | 14.48 | 0.19 |
| Post Translational Protein Modification | 25.90 | 0.08 |
| Pyruvate Metabolism and Citric Acid TCA Cycle | 0.98 | 0.05 |
| rRNA Processing | 0.00 | 0.00 |
| rRNA Processing in the Nucleus and Cytosol | 0.00 | 0.00 |
| Signaling by GPCR | 80.35 | 0.28 |
| Signaling by Interleukins | 50.68 | 0.29 |
| Signaling by Receptor Tyrosine Kinases | 56.88 | 0.34 |
| Signaling by Rho GTPases | 6.95 | 0.05 |
| Signaling by VEGF | 3.07 | 0.08 |
| Signaling by WNT | 9.53 | 0.10 |
| Transcriptional Regulation by TP53 | 4.83 | 0.08 |
| Translation | 1.02 | 0.03 |
| Transmission Across Chemical Synapses | 63.01 | 0.58 |
| Transport of Small Molecules | 24.13 | 0.11 |
| Vesicle Mediated Transport | 15.57 | 0.09 |

Many of the prioritised REACTOME pathways in Table 1 are associated with schizophrenia as reported in previous studies, such as signaling by GPCR [28, 32, 40], signaling by receptor tyrosine kinases [9, 17], signaling by interleukins [11, 33], and MAPK family signaling cascades [7, 15]. Although scored low by DiGSeE gene set scores, N-Linked glycosylation can play a modulatory role in neural transmission [38], and abnormalities in N-Linked glycosylation potentially indicate disruptions of central cell signaling processes in schizophrenia [29, 31]. Similarly, it has been reported that glucose oxidation, which takes place in the tricarboxylic acid (TCA) cycle, is inherently abnormal in a significant proportion of schizophrenic brains [5] and yet the corresponding REACTOME pathway only had low DiGSeE scores. Additional gene set prioritisation results corresponding to other evaluated GSCs are provided in Supplementary Results.

### Different network classifiers lead to different post-hoc interpretations

Functional edge selection results by different classifiers were better interpreted by different GSCs (Figure 2d) in the post-hoc interpretation framework. Edges selected by `glmnet` were best ex-plained by KEGG pathways, followed by the GO molecular function terms; see Supplementary Table S6 and Figure S7a. In contrast, edges selected by `graphClass` were better explained by GO molecular function terms, GO biological process terms and REACTOME pathways; see Supplementary Table S7 and Figure S7b.

As expected, GSCs with no obvious involvement in schizophrenia, such as the chromosomal position GSC and the immunological signature GSC, were not helpful in interpreting the functional edge selection results by any classifier. Figure 4 ranks the effectiveness of all examined GSCs by the AUC of each corresponding ROC curve for both classification methods. Generally, GO terms provided better interpretation compared to other GSCs by most of the metrics in both classification methods, and immunologic signatures and positional gene sets on chromosomes provided almost no insight in this case study.

**Figure 4:**
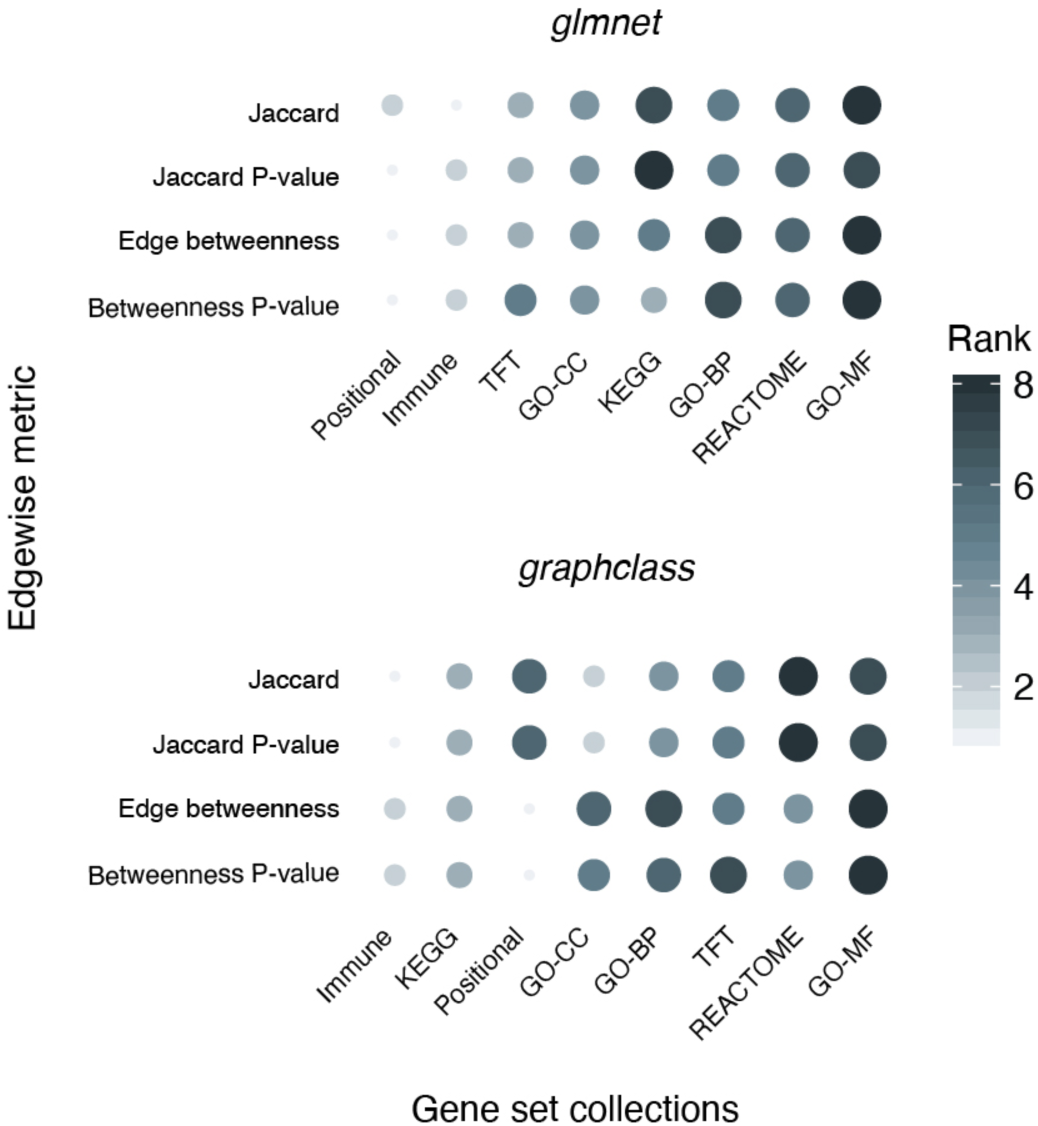
Interpretability of GSCs as measured by AUC in post-hoc interpretation framework for four different interpretation metrics (Jaccard, Jaccard *P*-value, Edge betweenness, and Edge betweenness *P*-value). ROC curves corresponding to each AUC value are depicted in Figure S7. Size and colour of the dots correspond to the rank of the GSC by AUC within each interpretation metric. For each classifier, GSCs on the horizontal axis are ordered by its average rank across all interpretation metrics. (AUC: area under the curve; ROC: receiver operating characteristic.)

## Discussion

We have devised a novel transcriptional-data-guided fMRI network classification algorithm via the strategic construction of feature groups in order to link transcriptional connectivity with functional connectivity across distinct brain regions with the aid of biologically meaningful gene set information. Application to publicly available data shows that our method provides better prediction accuracy in distinguishing fMRI networks of schizophrenic patients versus healthy controls while enabling biologically meaningful interpretation for selected functional connectivity features. In addition, we have also provided a post-hoc interpretation framework to enable biologically meaningful interpretation of functional edge selection results by other fMRI network classification algorithms.

Comparison among feature selection results guided by different GSCs of gene sets suggests there are potentially multiple functional connectivity biomarkers in schizophrenia. For example, two GSCs, GO molecular function terms and the REACTOME pathways, performed similarly well in terms of classification accuracy (Figure 3a) but had very different edge selection results (Supplementary Figure S6). This is by design, since different GSCs use different features and feature groups in training. Thus a biologically meaningful interpretation must consider selected functional connectivity features together with the corresponding GSC used in the construction of feature groups.

The choice of the GSC also affects the computational efficiency of the proposed classifier. Given a set of predictors, the number of feature groups, which depends on the number of gene sets in the selected GSC, increases the computational time of `brainClass` approximately exponentially. Another tuning parameter that affects the computational efficiency is the hard threshold on edge weights since a stricter cut-off reduces the number of features. An ad hoc investigation suggests that gene set rankings by the gene set prioritisation metric were relatively robust to the choice of the threshold used within a given GSC in our case study.

The ROC-based evaluation of the post-hoc interpretation framework substantially depends on how we construct disease association scores for each gene set. The scores were obtained by taking the sum or the average of association scores provided by DiGSeE for each individual gene after log-transformation, but the collective effect of a group of genes in disease mechanisms may not be additive. Besides, many studies have emphasised the role of pathways or biological processes in the pathogenesis of neurological disorders, rather than individual genes. For example, the WNT signalling pathway has been implicated in many neurodevelopment processes and was reported to be associated with schizophrenia [8, 21, 25], but none of these papers were counted towards the DiGSeE score because they do not mention any genes involved in the pathway within their abstracts. Similarly, this suggests that gene set prioritisation results will become more interpretable with the availability of better scoring systems for associations between diseases and molecular signatures such as pathways and GO terms.

One advantage of our framework is its flexibility. This approach can be readily applied for different GSCs, different brain parcellations, and different types of brain network data (e.g., diffusion weighted imaging). In addition, in the post-hoc interpretation framework, one can define different interpretation metrics. One could also utilise gene expression data in different ways to construct different types of feature groups for use in the `brainClass` method.

## Data Availability

Processed example data and the R implementation of all proposed methods are available at https://github.com/Mengbo-Li/brainClass.

## Acknowledgements

The authors thank all their colleagues at The University of Sydney, School of Mathematics and Statistics, for their support and intellectual engagement. We also thank Dr. Stephan F. Taylor and his research group at the University of Michigan for assistance with the data set.

## Author’s contributions

J.Y.H.Y., E.L., J.A., and D.K. conceived the study. M.L. developed the methods; performed data analysis and implemented the R package with input from J.A., D.K., E.L. and J.Y.H.Y.. J.Y.H.Y. and M.L. interpreted the results with input from J.A. and D.K. D.K., J.A., S.F. and M.B. processed and curated the data. All authors wrote, read and approved the final version of the manuscript.

## Supplementary Methods

### COBRE schizophrenia resting state fMRI data

Functional MRI (fMRI) networks for subjects from the Center for Biomedical Research Excellence (COBRE) schizophrenia resting state fMRI data (54 schizophrenic and 70 healthy control) [1, 8] were downloaded from the supplement to [2]. Details on the pre-processing of raw anatomic and functional scans from the COBRE data set are provided in [2].

### Allen Human Brain Atlas (AHBA) whole human brain gene expression microarray data

Brain-wide microarray gene expression data were obtained from the Allen Human Brain Atlas (AHBA) [4], which provides an anatomically comprehensive adult human brain transcriptome atlas comprising 3, 702 spatially distinct brain tissue samples obtained from six cadaver brains profiled for more than 20, 000 genes. During pre-processing, each sample was assessed for its quality and samples with poor quality spot plots, unusual plots of log-intensity ratios versus log-intensity averages or abnormal gene expression distributions were excluded, leaving 3, 546 samples. After pre-processing, the data were normalised using a conventional background correction in combination with quantile normalisation. Critically, all brain tissue samples were anatomically annotated to the normalised Montreal Neurological Institute (MNI) brain space (MNI152 non-linear). The MNI coordinates of each AHBA brain tissue sample were downloaded from https://github.com/chrisgorgo/alleninf/tree/master/alleninf/data.

### Power Brain Parcellation

Power et al. (2011) proposed a functional brain parcellation consisting of 264 anatomically distinct regions of interest (ROIs) comprising 14 functional brain systems modelled as 5*mm* radius spheres (Figure S1a) [7], whose centre coordinates were also mapped to the MNI152 1mm brain space. ROIs identified by the Power Parcellation were used as nodes to construct brain-wide fMRI networks and the gene set expression networks. In our analysis, pre-processed time series courses were extracted from voxels constituting each Power ROI from COBRE data, where node 75 is missing. The assignment of AHBA gene expression samples via sample MNI coordinates to the Power ROIs is described in the next section.

### Brain regional mapping of AHBA mircroarray samples to Power Parcellation ROIs

ROIs identified by the Power Parcellation [7] were used to construct gene set expression networks. To obtain gene expression profiles for each Power ROI, AHBA microarray samples were spatially mapped to the ROIs via their MNI coordinates. In order to obtain gene expression profiles for as many Power ROIs as possible, we enlarged the radius of spheres defined by the Power Parcellation to get enlarged ROIs for better coverage. The brain regional mapping from AHBA samples to (enlarged) Power ROIs has 3 steps (Figure S3):

**Step 1** Map the AHBA sample to a Power ROI if it lies within the 5*mm* radius from the centre of the ROI according to the original definition. Out of the 3, 546 preprocessed AHBA brain tissue samples, 240 samples got assigned to 118 ROIs.

**Step 2** Enlarge the radius of spheres defined as ROIs by the Power Parcellation [7]. While keeping the centre MRI coordinates of each sphere unchanged, we increase the radius of all spheres from 5*mm* to 12*mm*. Some ROIs become overlapped when *r* = 12*mm*. Note that samples can only be mapped into the non-overlapping regions of each ROI in this step. The enlarged radius of 12*mm* was picked so that as many AHBA samples were mapped to the non-overlapped regions as possible (Figure S2). 1, 129 gene expression samples were mapped to 196 different enlarged ROIs in this step. So far, 1, 369 AHBA samples were mapped to 211 enlarged ROIs.

**Step 3** Map the remaining AHBA gene expression samples to the so-far empty enlarged ROIs (*r* = 12*mm*). In this step, 119 samples were mapped to 52 enlarged ROIs.

Eventually after all three steps, 1, 488 AHBA microarray samples were mapped into 248 enlarged Power ROIs.

### Dimension reduction on the AHBA microarray data

After the initial data pre-processing and normalisation on the AHBA gene expression data, we obtained a 3, 546 × 32, 488 data matrix, where each row corresponds to an AHBA brain tissue sample and each column corresponds to a probe ID. We first filtered out probes that were not annotated to any HGNC gene symbols and tissue samples that were not assigned to any ROI. This returned us a sample ID by gene symbol data matrix (1, 488 × 19, 227). We discarded ROI75 as it is missing in COBRE data. For each gene, we calculated its average expression level in samples assigned to the same Power ROI to obtain a ROI by gene symbol data matrix (247 × 19, 227). We further filtered out genes with low expression in all ROIs. Average gene expression levels across 247 ROIs for each of the 19, 227 genes were calculated and those below the 3rd quartile were filtered out. We ended up with a 247 × 4, 807 ROI by gene symbol data matrix, where each entry in the matrix contains the average expression level of a gene within the enlarged Power ROI (Figure S1b).

### Molecular Signatures Database (MSigDB, v7.0)

Gene set collections from the Molecular Signatures Database (MSigDB, v7.0) are publicly available at http://software.broadinstitute.org/gsea/msigdb/index.jsp, which is a comprehensive gene set database that is commonly used to perform gene set enrichment analysis (GSEA), providing labelled gene set information on biological processes and disease pathways to enable biologically meaningful interpretation of transcriptional data [6]. Typically, each category of gene set collections represent a particular domain of knowledge [6]. For instance, gene ontology (GO) [3] gene sets are derived based on hierarchically controlled GO terms including molecular function (MF), cellular component (CC) or biological process (BP) to provide meaningful annotations for gene functions and products. Alternatively, gene sets can also be curated simply by the physical location of each gene on the chromosomes (the positional gene set collection). MSigDB also provides processed and annotated gene sets that constitute various types of biological pathways such as KEGG or REACTOME.

### Gene set schizophrenia association scores based on DiGSeE

Disease gene search engine with evidence sentences (DiGSeE) [5] is a web tool for gene-disease association queries (http://gcancer.org/digsee). The database searches for genes associated with the disease of interest (e.g. schizophrenia) in current literature, and provides a score for each gene, based on joint occurrences of the disease and relevant genes through biological events in MEDLINE abstracts. Gene-schizophrenia association scores were obtained from DiGSeE for our case study. To account for different aliases of the same gene, we summed association scores for gene symbols mapped to the same entrez ID. A natural log-transformation was applied to raw scores due to its right-skewed distribution. We then filtered out genes with a relatively low schizophrenia association score by keeping only genes scored higher than or equal to the median of log-transformed scores.

After pre-processing, we constructed two types of schizophrenia relevance scores for each gene set: *DiGSeE sum score* and *DiGSeE average score*. We first filtered each gene set by removing genes previously identified as having low expression in gene expression data and discarded gene sets consisting of less than 5 genes post filtering. Gene set schizophrenia relevance scores were calculated by taking the sum and the average of gene-schizophrenia association scores with genes from each filtered gene set (Figure S1c).

### Receiver operating characteristic (ROC) curve and area under the curve (AUC)

Within each gene set collection (GSC), gene sets were ranked by post-hoc interpretability metrics. True positives (gene sets associated with schizophrenia) and true negatives were defined by summarised gene set schizophrenia relevance scores based on DiGSeE. By each interpretability metric, gene sets that are ranked top *x* percent (from 0% to 100% at a step of 5%) were categorised as test positives (associated with schizophrenia) and test negatives (not associated with schizophrenia) otherwise. The accuracy of gene set categorisation by each metric was evaluated by the area under the receiver operating characteristic (ROC) curve, where the true positive rate is plotted against the false positive rate by comparing true positives with test positives. Complete results are provided in Figure S7, with AUC for each curve in the figure provided in Table S6 and Table S7.

## Supplementary Results

### Gene set prioritisation results when guided by different collections of gene sets

Given selected functional edge features by the classifier, a gene set prioritisation metric was calculated as described in the Methods section. Gene sets were then ranked within each GSC by the prioritisation metric. Gene sets ranked top 5% within each GSC in all ten folds of all 50 repetitons of 10-fold cross-validations are provided in Tables S1 to S5. For each GSC, distributions of gene set schizophrenia relevance scores for (i) all gene sets in each GSC and (ii) the prioritised gene sets are provided in Figure S4.

### Number of selected features when guided by different GSCs

Given a particular GSC, we observed that the distribution of the number of selected features (around 10% of all included features are selected) and the prediction accuracy are robust to the regularisation parameter λ in `grpregOverlap`, when 1 × 10^−3^ ≤ λ < 1 × 10^−2^. In our implementation, the default is λ = 5 × 10^−3^. The number of features included in classification when guided by different GSCs, and the median number of selected features (that had non-zero coefficients) in 10-fold cross-validations over 50 repeats are summarised in Figure S5.

## Supplementary Figures

**Figure S1:**
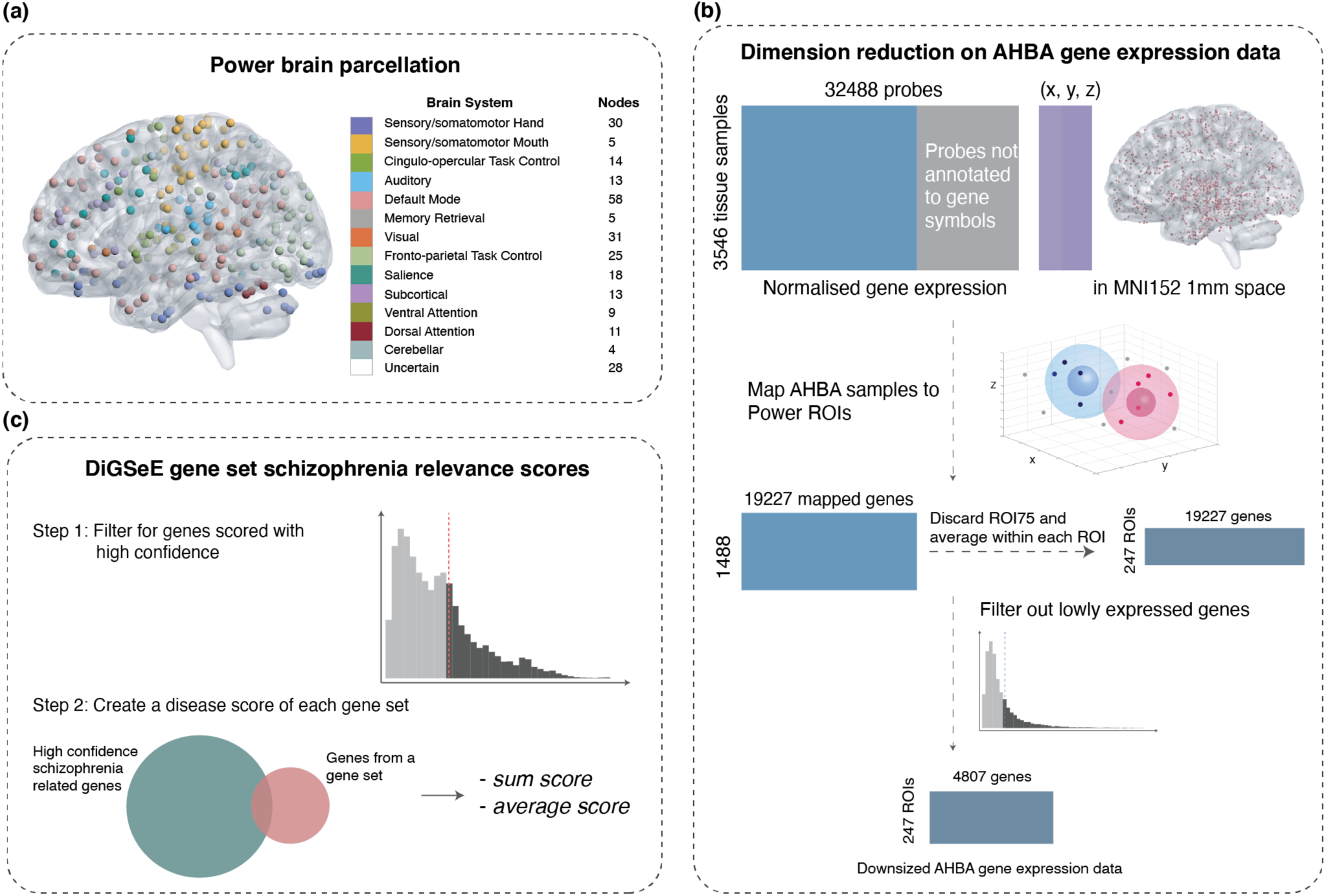
Data pre-processing. (a) The Power brain parcellation. Regions of interest (ROIs) are modelled as 5*mm* radius spheres and coloured by functional brain systems. The total number of ROIs in each system are also indicated. This illustration was created using the BrainNet Viewer Matlab software [9]. (b) Dimension reduction on the Allen Human Brain Atlas (AHBA) gene expression data. (c) Construction of schizophrenia relevance scores for each gene set based on the gene-disease association query results from DiGSeE.

**Figure S2:**
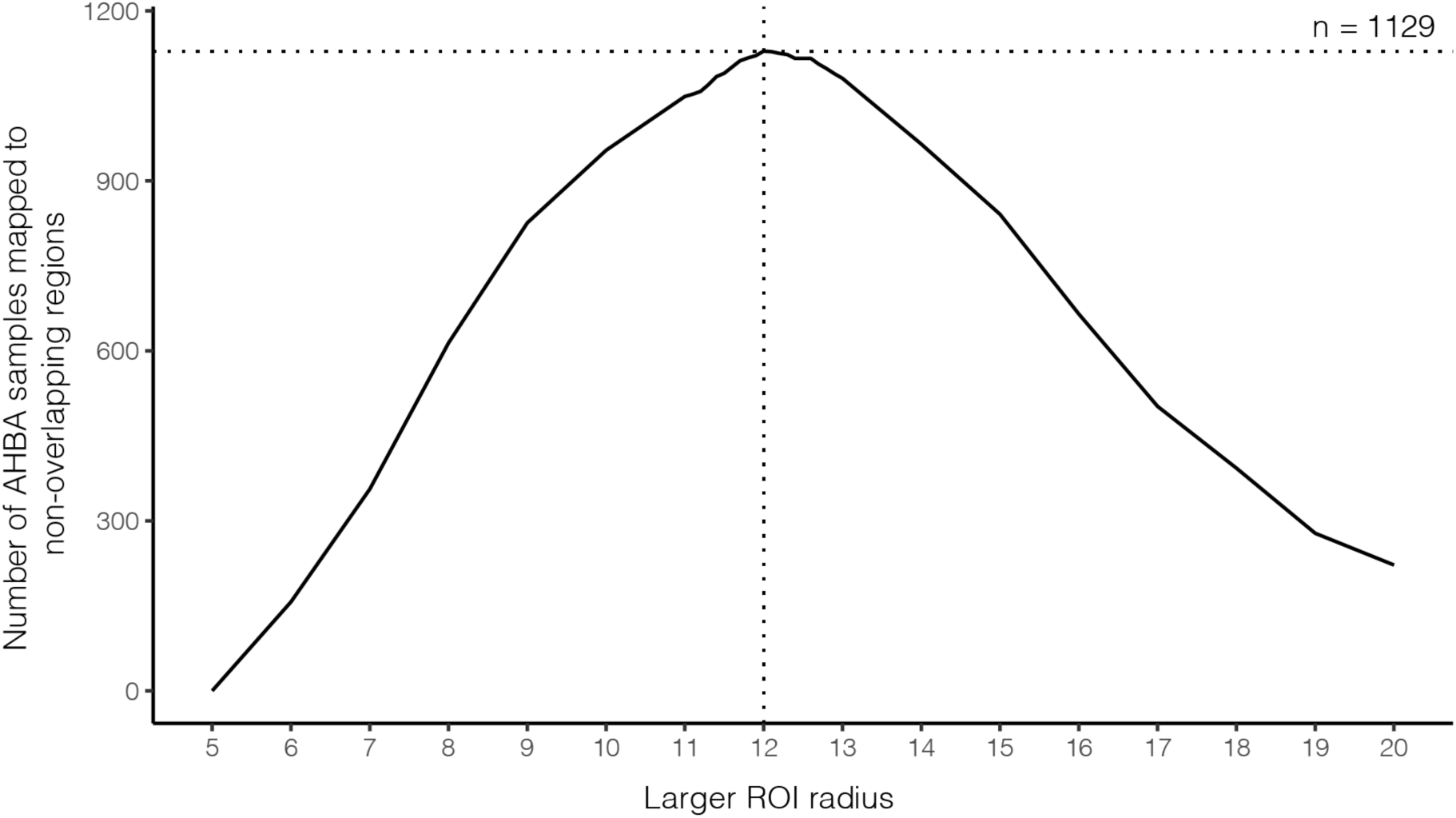
Number of AHBA samples mapped to the non-overlapping regions of enlarged ROIs as we increase the radius of Power ROI spheres. Originally, Power ROIs are modelled as *r* = 5*mm* spheres, and none of them overlap. In order to obtain gene expression profiles for as many ROIs as possible, we increased the radius of all spheres at a step of 0.01*mm*, from 5*mm* to 20*mm*. As the radius of all ROIs increase, some of them start to overlap, and we only map the AHBA samples located in the non-overlapping regions to its closest ROI.

**Figure S3:**
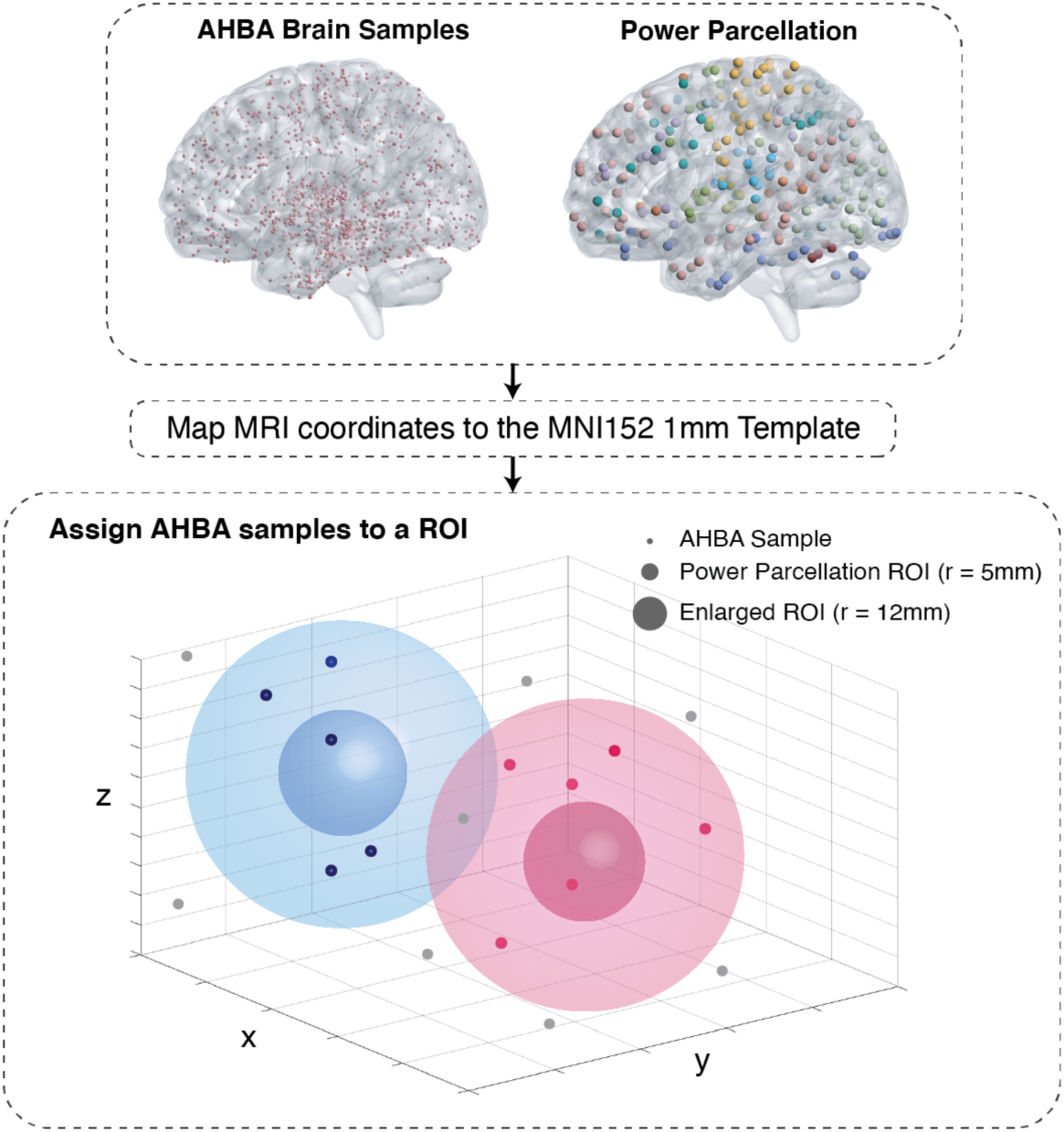
Schematic illustration of the AHBA microarray sample assignment to the Power brain parcellation.

**Figure S4:**
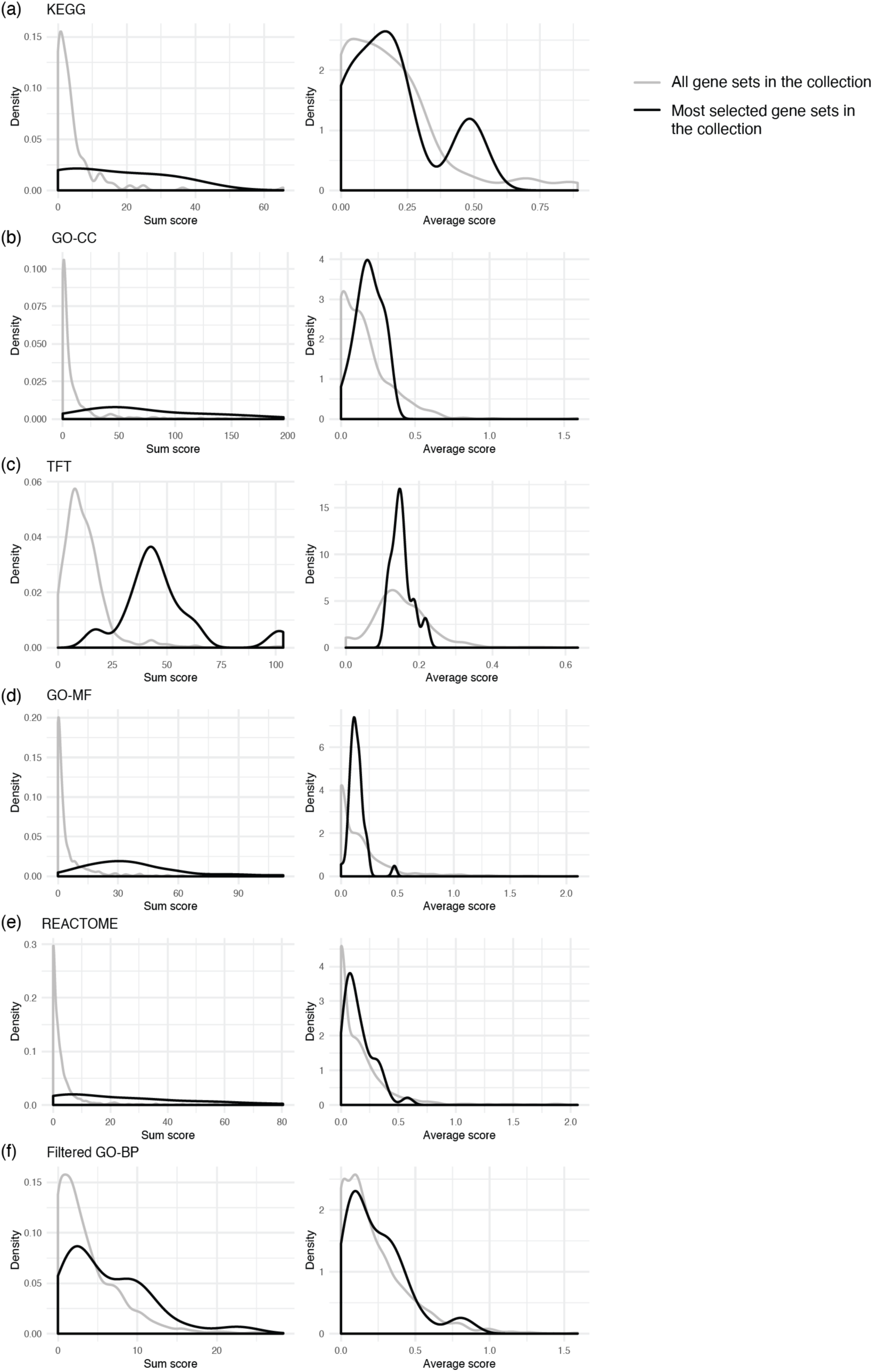
Gene set schizophrenia association score distributions. All gene set collections were filtered and processed as described in the Supplementary Methods section. Distributions of all gene set schizophrenia relevance scores are plotted in grey. Plotted in black are the distributions of the gene set association scores for those selected in all 50 repeats of 10-fold cross-validations in the case study. The filtered GO biological process (BP) term collection was obtained by only keeping those sized between 10 and 30. (Abbreviations: TFT: transcription factor target; CC: cellular component; MF: molecular function.)

**Figure S5:**
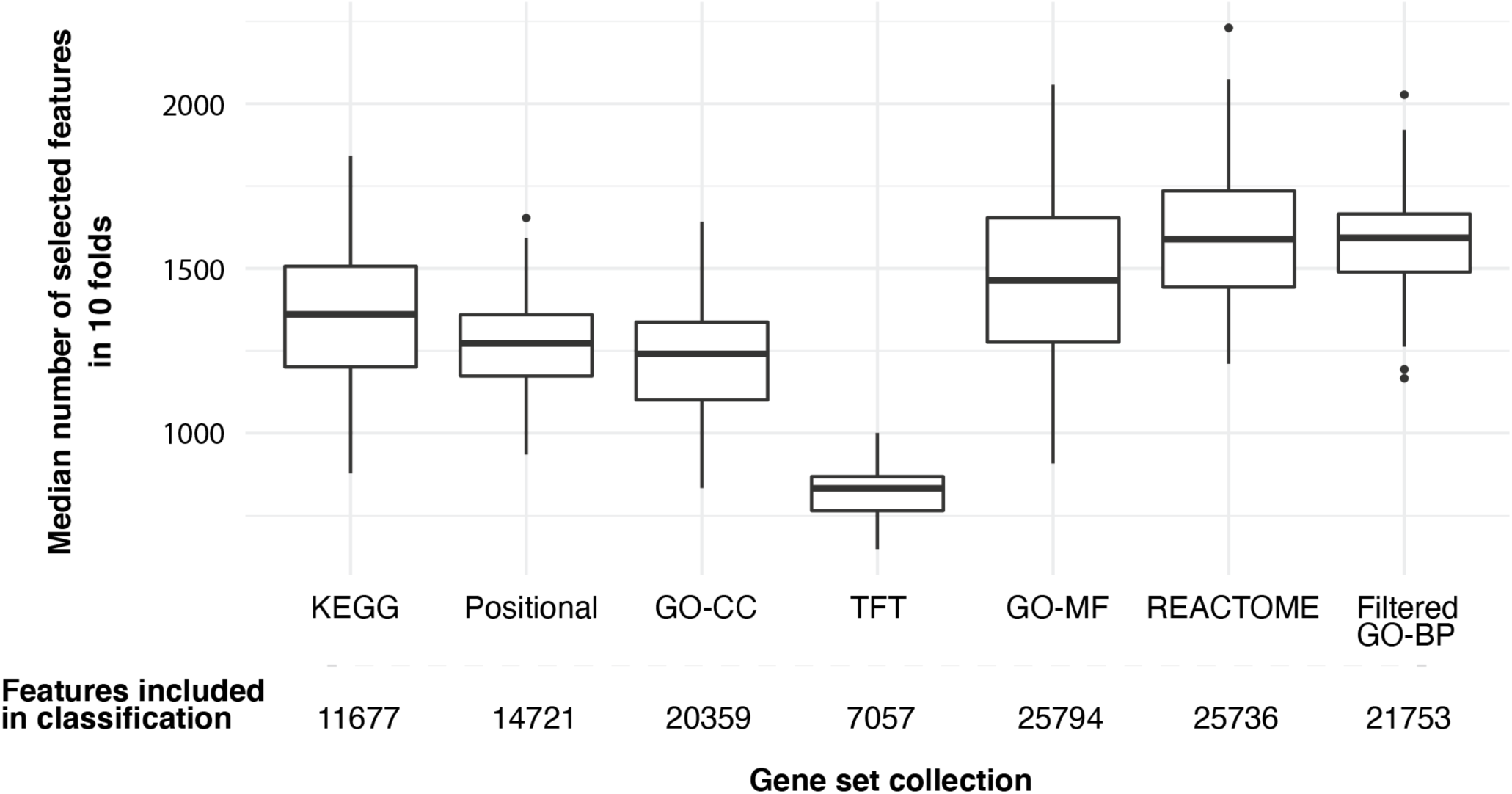
Median number of selected features in 10-fold cross-validations over 50 trials by the gene expression guided classification algorithm (λ= 5 × 10^−3^). By design, edges that do not belong to any gene set edge groups are excluded from model training, thus the selection of GSC directly determines which features are included in classification. The numbers of features included in model training are provided.

**Figure S6:**
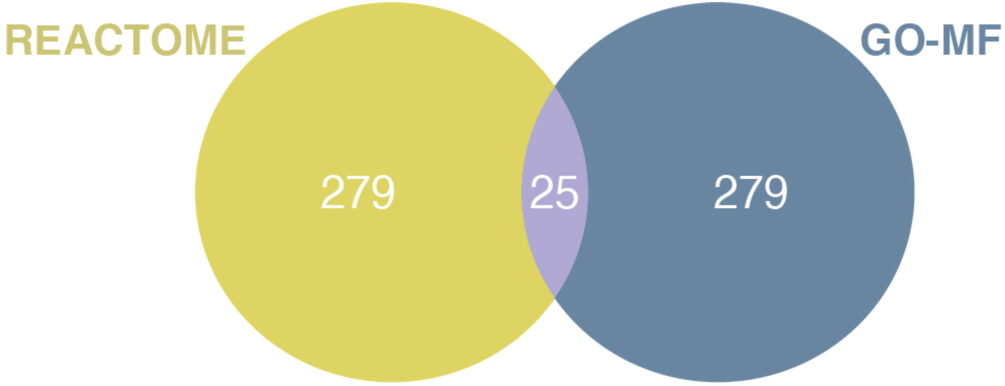
The overlap between most selected functional edges when the proposed classifier was guided by REACTOME pathways versus the gene ontology (GO) molecular function (MF) terms. Most selected edges were edges selected in all 10 folds and all 50 trials of cross-validations.

**Figure S7:**
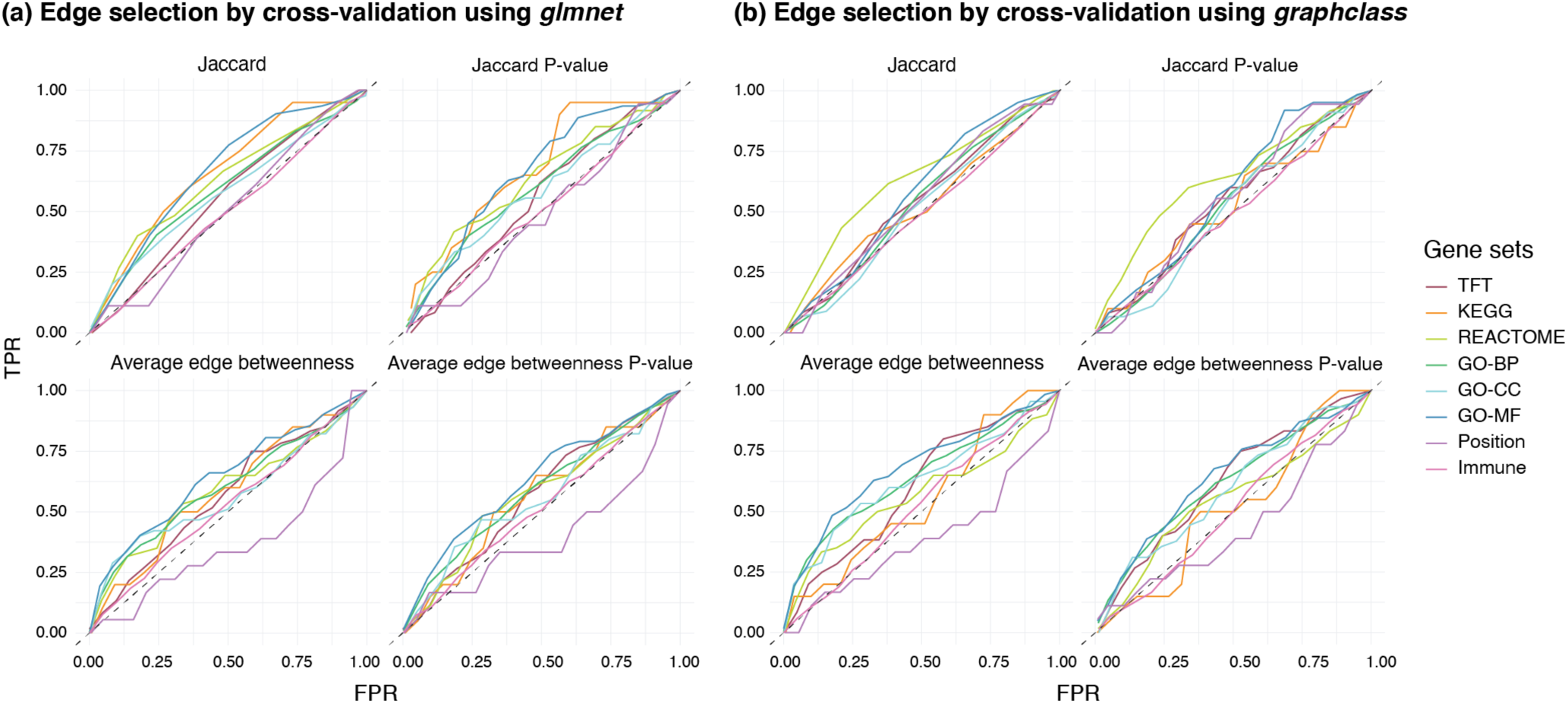
Receiver operating characteristic (ROC) curve evaluation of each post-hoc interpretability metric when different GSCs were used for interpreting predictive functional edges selected by (a) `glmnet` and (b) `graphclass`. (FPR: false positive rate; TPR: true positive rate.)

## Supplementary Tables

**Table S1:** KEGG pathway gene sets with a prioritisation score in top 5% in all ten folds in all 50 repeats of 10-fold cross-validations in alphabetical order.

| Gene set | Sum score | Average score |
| --- | --- | --- |
| Calcium Signaling Pathway | 36.32 | 0.48 |
| Citrate Cycle TCA Cycle | 0.00 | 0.00 |
| Glycolysis Gluconeogenesis | 0.96 | 0.09 |
| Pathways in Cancer | 25.22 | 0.23 |
| Regulation of Actin Cytoskeleton | 12.18 | 0.17 |

**Table S2:** Transcription factor target gene sets with a prioritisation score in top 5% in all ten folds in all 50 repeats of 10-fold cross-validations in alphabetical order.

| Gene set | Sum score | Average score |
| --- | --- | --- |
| AAANWWTGC_UNKNOWN | 16.69 | 0.19 |
| AAAYRNCTG_UNKNOWN | 28.68 | 0.22 |
| AACTTT_UNKNOWN | 103.36 | 0.14 |
| CACGTG_MYC_Q2 | 41.78 | 0.14 |
| CAGCTG_AP4_Q5 | 58.25 | 0.11 |
| CAGGTA_AREB6_01 | 42.10 | 0.15 |
| CTGCAGY_UNKNOWN | 63.28 | 0.22 |
| CTTTAAR_UNKNOWN | 48.13 | 0.14 |
| CTTTGA_LEF1_Q2 | 54.82 | 0.14 |
| GATTGGY_NFY_Q6_01 | 37.46 | 0.12 |
| GCANCTGNY_MYOD_Q6 | 51.12 | 0.15 |
| GGGTGGRR_PAX4_03 | 51.60 | 0.12 |
| RNGTGGGC_UNKNOWN | 43.77 | 0.15 |
| RTAAACA_FREAC2_01 | 39.18 | 0.12 |
| RYTTCCTG_ETS2_B | 43.62 | 0.13 |
| TAATTA_CHX10_01 | 43.57 | 0.14 |
| TATAAA_TATA_01 | 63.05 | 0.16 |
| TCANNTGAY_SREBP1_01 | 17.32 | 0.12 |
| TGACAGNY_MEIS1_01 | 42.53 | 0.15 |
| TGACATY_UNKNOWN | 36.29 | 0.15 |
| TGACCTY_ERR1_Q2 | 47.17 | 0.13 |
| TGCCAAR_NF1_Q6 | 45.08 | 0.17 |
| TGGAAA_NFAT_Q4_01 | 99.81 | 0.16 |
| TGTTTGY_HNF3_Q6 | 40.36 | 0.16 |
| WGTTNNNNNAAA_UNKNOWN | 33.75 | 0.19 |
| WTTGKCTG_UNKNOWN | 33.38 | 0.19 |

**Table S3:** Gene ontology cellular component gene sets with a prioritisation score in top 5% in all ten folds in all 50 repeats of 10-fold cross-validations in alphabetical order.

| Gene set | Sum score | Average score |
| --- | --- | --- |
| Axon | 82.88 | 0.32 |
| Catalytic Complex | 9.51 | 0.04 |
| Cell Body | 51.17 | 0.23 |
| Cell Junction | 89.17 | 0.17 |
| Cell Projection Part | 123.33 | 0.24 |
| Cytoskeletal Part | 40.62 | 0.09 |
| Endoplasmic Reticulum | 66.17 | 0.13 |
| Envelope | 37.04 | 0.15 |
| Filopodium | 9.84 | 0.20 |
| Golgi Apparatus | 42.91 | 0.10 |
| Golgi Apparatus Part | 20.93 | 0.08 |
| Intrinsic Component of Plasma Membrane | 137.94 | 0.25 |
| Membrane Protein Complex | 60.51 | 0.20 |
| Mitochondrion | 46.46 | 0.14 |
| Neuron Part | 196.16 | 0.28 |
| Neuron Projection | 159.93 | 0.31 |
| Nuclear Chromosome Telomeric Region | 0.00 | 0.00 |
| Perinuclear Region of Cytoplasm | 45.77 | 0.19 |
| Plasma Membrane Region | 121.90 | 0.27 |
| Somatodendritic Compartment | 107.55 | 0.31 |
| Supramolecular Complex | 39.87 | 0.15 |
| Synapse | 161.38 | 0.31 |
| Vesicle Membrane | 46.68 | 0.20 |
| Whole Membrane | 79.23 | 0.16 |

**Table S4:**
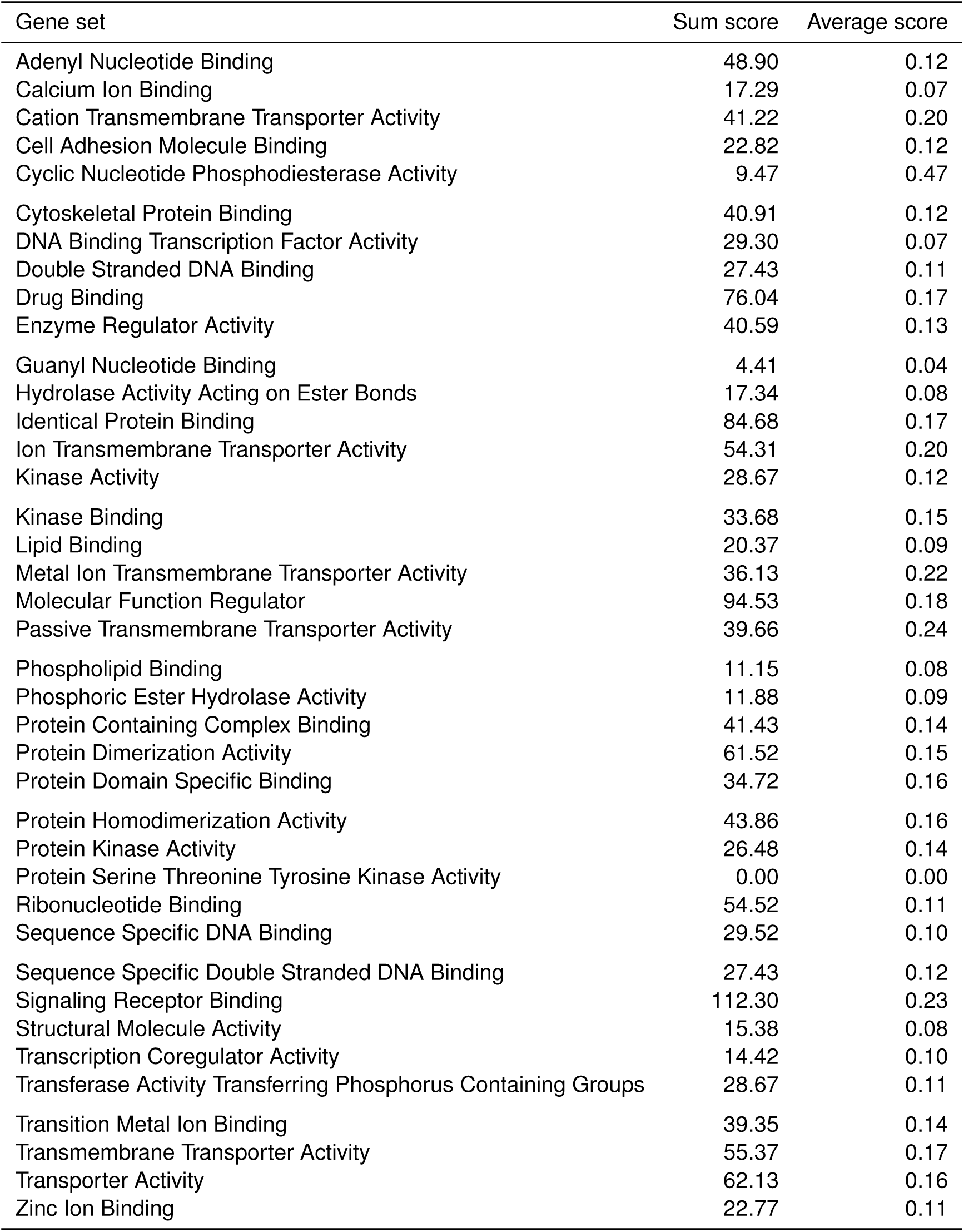
Gene ontology molecular function gene sets with a prioritisation score in top 5% in all ten folds in all 50 repeats of 10-fold cross-validations in alphabetical order.

| Gene set | Sum score | Average score |
| --- | --- | --- |
| Adenyl Nucleotide Binding | 48.90 | 0.12 |
| Calcium Ion Binding | 17.29 | 0.07 |
| Cation Transmembrane Transporter Activity | 41.22 | 0.20 |
| Cell Adhesion Molecule Binding | 22.82 | 0.12 |
| Cyclic Nucleotide Phosphodiesterase Activity | 9.47 | 0.47 |
| Cytoskeletal Protein Binding | 40.91 | 0.12 |
| DNA Binding Transcription Factor Activity | 29.30 | 0.07 |
| Double Stranded DNA Binding | 27.43 | 0.11 |
| Drug Binding | 76.04 | 0.17 |
| Enzyme Regulator Activity | 40.59 | 0.13 |
| Guanyl Nucleotide Binding | 4.41 | 0.04 |
| Hydrolase Activity Acting on Ester Bonds | 17.34 | 0.08 |
| Identical Protein Binding | 84.68 | 0.17 |
| Ion Transmembrane Transporter Activity | 54.31 | 0.20 |
| Kinase Activity | 28.67 | 0.12 |
| Kinase Binding | 33.68 | 0.15 |
| Lipid Binding | 20.37 | 0.09 |
| Metal Ion Transmembrane Transporter Activity | 36.13 | 0.22 |
| Molecular Function Regulator | 94.53 | 0.18 |
| Passive Transmembrane Transporter Activity | 39.66 | 0.24 |
| Phospholipid Binding | 11.15 | 0.08 |
| Phosphoric Ester Hydrolase Activity | 11.88 | 0.09 |
| Protein Containing Complex Binding | 41.43 | 0.14 |
| Protein Dimerization Activity | 61.52 | 0.15 |
| Protein Domain Specific Binding | 34.72 | 0.16 |
| Protein Homodimerization Activity | 43.86 | 0.16 |
| Protein Kinase Activity | 26.48 | 0.14 |
| Protein Serine Threonine Tyrosine Kinase Activity | 0.00 | 0.00 |
| Ribonucleotide Binding | 54.52 | 0.11 |
| Sequence Specific DNA Binding | 29.52 | 0.10 |
| Sequence Specific Double Stranded DNA Binding | 27.43 | 0.12 |
| Signaling Receptor Binding | 112.30 | 0.23 |
| Structural Molecule Activity | 15.38 | 0.08 |
| Transcription Coregulator Activity | 14.42 | 0.10 |
| Transferase Activity Transferring Phosphorus Containing Groups | 28.67 | 0.11 |
| Transition Metal Ion Binding | 39.35 | 0.14 |
| Transmembrane Transporter Activity | 55.37 | 0.17 |
| Transporter Activity | 62.13 | 0.16 |
| Zinc Ion Binding | 22.77 | 0.11 |

**Table S5:** Gene ontology biological process gene sets with a prioritisation score in top 5% in all ten folds in all 50 repeats of 10-fold cross-validations in alphabetical order.

| Gene set | Sum score | Average score |
| --- | --- | --- |
| Acute Inflammatory Response | 10.40 | 0.36 |
| Amyloid Precursor Protein Metabolic Process | 10.90 | 0.44 |
| Artery Development | 1.99 | 0.07 |
| Axis Specification | 2.10 | 0.08 |
| Catecholamine Secretion | 15.76 | 0.58 |
| Cell Adhesion Mediated by Integrin | 0.94 | 0.03 |
| Cell Differentiation Involved in Kidney Development | 2.09 | 0.10 |
| Cellular Response to Heat | 6.69 | 0.22 |
| Cellular Response to Lipoprotein Particle Stimulus | 2.97 | 0.30 |
| Cerebral Cortex Cell Migration | 9.61 | 0.42 |
| Diencephalon Development | 9.94 | 0.37 |
| DNA Damage Response Signal Transduction by P53 Class Mediator | 2.57 | 0.10 |
| Embryonic Heart Tube Development | 0.00 | 0.00 |
| Embryonic Pattern Specification | 6.79 | 0.27 |
| Epithelial Cell Apoptotic Process | 1.64 | 0.06 |
| Fibroblast Growth Factor Receptor Signaling Pathway | 5.72 | 0.21 |
| Forebrain Generation of Neurons | 12.53 | 0.42 |
| Forebrain Neuron Differentiation | 11.31 | 0.44 |
| Glomerulus Development | 1.28 | 0.05 |
| Inner Ear Morphogenesis | 5.27 | 0.18 |
| Multi Organism Behavior | 13.45 | 0.50 |
| Muscle Cell Migration | 2.60 | 0.11 |
| Myeloid Cell Development | 6.77 | 0.26 |
| Negative Regulation of Extrinsic Apoptotic Signaling Pathway | 8.95 | 0.30 |
| Negative Regulation of Fat Cell Differentiation | 1.02 | 0.06 |
| Negative Regulation of Hormone Secretion | 9.61 | 0.34 |
| Negative Regulation of Ossification | 0.87 | 0.03 |
| Negative Regulation of Protein Polymerization | 3.24 | 0.12 |
| Neural Crest Cell Migration | 8.85 | 0.34 |
| Neuroblast Proliferation | 11.05 | 0.38 |
| Neuron Fate Commitment | 8.40 | 0.32 |
| Neuron Recognition | 7.99 | 0.27 |
| Odontogenesis of Dentin Containing Tooth | 0.00 | 0.00 |
| Osteoclast Differentiation | 0.87 | 0.03 |
| Pigmentation | 9.25 | 0.32 |
| Platelet Derived Growth Factor Receptor Signaling Pathway | 0.96 | 0.06 |
| Positive Regulation of Actin Filament Bundle Assembly | 2.03 | 0.08 |
| Positive Regulation of Animal Organ Morphogenesis | 5.30 | 0.20 |
| Positive Regulation of Cell Cycle Phase Transition | 2.05 | 0.09 |
| Positive Regulation of DNA Metabolic Process | 0.00 | 0.00 |
| Positive Regulation of Mitotic Cell Cycle | 8.97 | 0.31 |
| Positive Regulation of Muscle Tissue Development | 17.21 | 0.82 |
| Positive Regulation of Peptide Hormone Secretion | 3.90 | 0.14 |
| Positive Regulation of Protein Binding | 11.11 | 0.38 |
| Positive Regulation of Regulated Secretory Pathway | 2.91 | 0.13 |

Table S5: (Continued) Gene ontology biological process gene sets with a prioritisation score in top 5% in all ten folds in all 50 repeats of 10-fold cross-validations in alphabetical order.
| Gene set | Sum score | Average score |
| --- | --- | --- |
| Post Embryonic Development | 8.61 | 0.30 |
| Proteoglycan Biosynthetic Process | 2.95 | 0.11 |
| Regulation of Adherens Junction Organization | 0.96 | 0.03 |
| Regulation of Cardiac Muscle Tissue Development | 21.31 | 0.79 |
| Regulation of Cartilage Development | 1.06 | 0.04 |
| Regulation of Cellular Ketone Metabolic Process | 4.37 | 0.15 |
| Regulation of Chondrocyte Differentiation | 1.06 | 0.05 |
| Regulation of Filopodium Assembly | 3.58 | 0.14 |
| Regulation of Jun Kinase Activity | 4.80 | 0.19 |
| Regulation of Kidney Development | 3.37 | 0.18 |
| Regulation of Muscle Adaptation | 8.83 | 0.29 |
| Regulation of Neuronal Synaptic Plasticity | 11.04 | 0.39 |
| Regulation of Protein Targeting | 3.29 | 0.12 |
| Regulation of Protein Tyrosine Kinase Activity | 22.48 | 0.75 |
| Regulation of Smooth Muscle Contraction | 8.19 | 0.27 |
| Regulation of Stem Cell Proliferation | 5.20 | 0.19 |
| Response to Amino Acid | 11.30 | 0.45 |
| Response to Anesthetic | 24.53 | 0.88 |
| Response to Nerve Growth Factor | 13.77 | 0.55 |
| Response to Vitamin | 3.06 | 0.11 |
| Signal Transduction in Absence of Ligand | 6.55 | 0.25 |
| Signal Transduction in Response to DNA Damage | 2.57 | 0.09 |
| Smooth Muscle Cell Migration | 2.60 | 0.14 |
| Sterol Transport | 2.08 | 0.08 |
| Ventricular Septum Development | 1.93 | 0.07 |

**Table S6:** AUC of the corresponding ROC curve when interpreting *glmnet* functional edge selection results by each GSC with each post-hoc interpretability metric.

| Gene set collection | Jaccard | Jaccard P-value | Edge betweenness | Edge betweenness P-value |
| --- | --- | --- | --- | --- |
| TFT | 0.56 | 0.55 | 0.56 | 0.57 |
| KEGG | 0.66 | 0.67 | 0.58 | 0.56 |
| REACTOME | 0.63 | 0.63 | 0.59 | 0.57 |
| GO-BP | 0.59 | 0.59 | 0.60 | 0.59 |
| GO-CC | 0.58 | 0.58 | 0.57 | 0.56 |
| GO-MF | 0.66 | 0.66 | 0.64 | 0.63 |
| Positional | 0.51 | 0.48 | 0.37 | 0.40 |
| Immunological | 0.50 | 0.50 | 0.52 | 0.52 |

**Table S7:** AUC of the corresponding ROC curve when interpreting *graphclass* functional edge selection results by each GSC with each post-hoc interpretability metric.

| Gene set collection | Jaccard | Jaccard P-value | Edge betweenness | Edge betweenness P-value |
| --- | --- | --- | --- | --- |
| TFT | 0.54 | 0.55 | 0.60 | 0.62 |
| KEGG | 0.52 | 0.53 | 0.55 | 0.52 |
| REACTOME | 0.63 | 0.64 | 0.57 | 0.55 |
| GO-BP | 0.53 | 0.53 | 0.64 | 0.62 |
| GO-CC | 0.51 | 0.51 | 0.63 | 0.59 |
| GO-MF | 0.58 | 0.58 | 0.68 | 0.64 |
| Positional | 0.54 | 0.56 | 0.39 | 0.44 |
| Immunological | 0.49 | 0.49 | 0.52 | 0.50 |

